# Peripheral neuropathy linked mRNA export factor GANP reshapes gene regulation in human motor neurons

**DOI:** 10.1101/2021.05.18.444636

**Authors:** Rosa Woldegebriel, Jouni Kvist, Matthew White, Matilda Sinkko, Satu Hänninen, Markus T Sainio, Rubén Torregrosa-Munumer, Sandra Harjuhaahto, Nadine Huber, Sanna-Kaisa Herukka, Annakaisa Haapasalo, Olli Carpen, Andrew Bassett, Emil Ylikallio, Jemeen Sreedharan, Henna Tyynismaa

## Abstract

Loss-of-function of the mRNA export protein GANP (*MCM3AP* gene) cause early-onset sensorimotor neuropathy, characterised by axonal degeneration in long peripheral nerves. GANP functions as a scaffold at nuclear pore complexes, contributing to selective nuclear export of mRNAs. Here, we aimed to identify motor neuron specific transcripts that are regulated by GANP and may be limiting for local protein synthesis in motor neuron axons. We compared motor neurons with a gene edited mutation in the Sac3 mRNA binding domain of GANP to isogenic controls. We also examined patient-derived motor neurons. RNA sequencing of motor neurons as well as nuclear and axonal subcompartments showed that mutant GANP had a profound effect on motor neuron transcriptomes, with alterations in nearly 40 percent of all expressed genes and broad changes in splicing. Expression changes in multiple genes critical for neuronal functions, combined with compensatory upregulation of protein synthesis and early-stage metabolic stress genes, indicated that RNA metabolism was abnormal in GANP-deficient motor neurons. Surprisingly, limited evidence was found for large-scale nuclear retention of mRNA. This first study of neuropathy-linked GANP defects in human motor neurons shows that GANP has a wide gene regulatory role in a disease-relevant cell type that requires long-distance mRNA transport.

## INTRODUCTION

Gene expression - from mRNA synthesis to splicing, maturation and export from the nucleus to the cytoplasm for translation - is an interconnected process mediated through large macromolecular complexes [13,31,51]. For example, the nuclear pore complex (NPC), which is a large protein assembly embedded in the nuclear membrane, is not merely a gateway enabling nucleocytoplasmic transport but also affects upstream gene expression regulation [17,54]. Associating with the nuclear basket of the NPC, germinal centre associated nuclear protein (GANP) is involved in mRNA export acting as a large scaffold in the TRanscription and EXport-2 complex (TREX-2) [20,59]. GANP, a 210 kDa protein, has N-terminal FG (phenylalanine/glycine) repeats, a yeast Sac3 mRNA binding homology domain, and a C-terminal acetyltransferase domain. GANP has been described as a mediator of selective mRNA export, thus facilitating dynamic regulation of specific biological processes [57,58].

We and others recently identified GANP to underlie a childhood-onset neurological syndrome, caused by biallelic mutations in its encoding gene *MCM3AP* (MIM #618124, PNRIID) [23,24,49,61,63]. The disease affects peripheral nerves causing Charcot-Marie-Tooth neuropathy, and results in gait difficulties, often with loss of ambulation. Many patients have intellectual disability. Variable other central nervous system involvement such as encephalopathy or epilepsy have been reported in some patients, as well as non-neurological symptoms such as ovarian dysfunction [61,63]. Studies in patient fibroblasts showed that pathological *MCM3AP* variants either result in a decrease in the amount of GANP at the nuclear envelope or affect critical amino acids in the Sac3 mRNA binding domain of GANP [61]. Depletion of GANP was associated with more severe motor phenotypes than the Sac3 variants.

Our identification of GANP loss-of-function as a cause for human neurological disease suggested a key role for GANP in neuronal cells, and particularly in motor neurons. Interestingly, GANP was previously found to modify TDP-43 toxicity in fly motor neurons [50]. However, the functions of GANP, and its roles in gene regulation and mRNA export selectivity in human neurons remain unresolved. We hypothesised that inefficient mRNA export limiting the availability of mRNAs for local protein synthesis in axon terminals could be a pathogenic mechanism in GANP-associated disease. Here we set out to investigate the effects of GANP variants in human induced pluripotent stem cell (hiPSC) derived motor neurons. Our results show that GANP mutant motor neurons differentiate normally in culture but have altered expression of nearly 40 percent of their genes, and major changes in gene splicing. Thus, we suggest that GANP has a key gene regulatory role in human motor neurons beyond scaffolding for mRNA export.

## RESULTS

### *MCM3AP* gene edited and patient-derived hiPSC are pluripotent

We used CRISPR-Cas9 gene editing to knock in different *MCM3AP* patient mutations in the kolf2_c1 parental hiPSC line [26] by homology-directed repair [3]. 128 clones were screened for mutations by MiSeq, and successful editing was detected for the patient mutation R878H located in the Sac3 domain of GANP [23] (Figure 1A,B). Sanger sequencing of the edited hiPSC clones showed that one clone was homozygous for the mutation. Karyotypes of the homozygous edited and parental lines were as expected 46 XY by g-banding.

**Figure 1.**
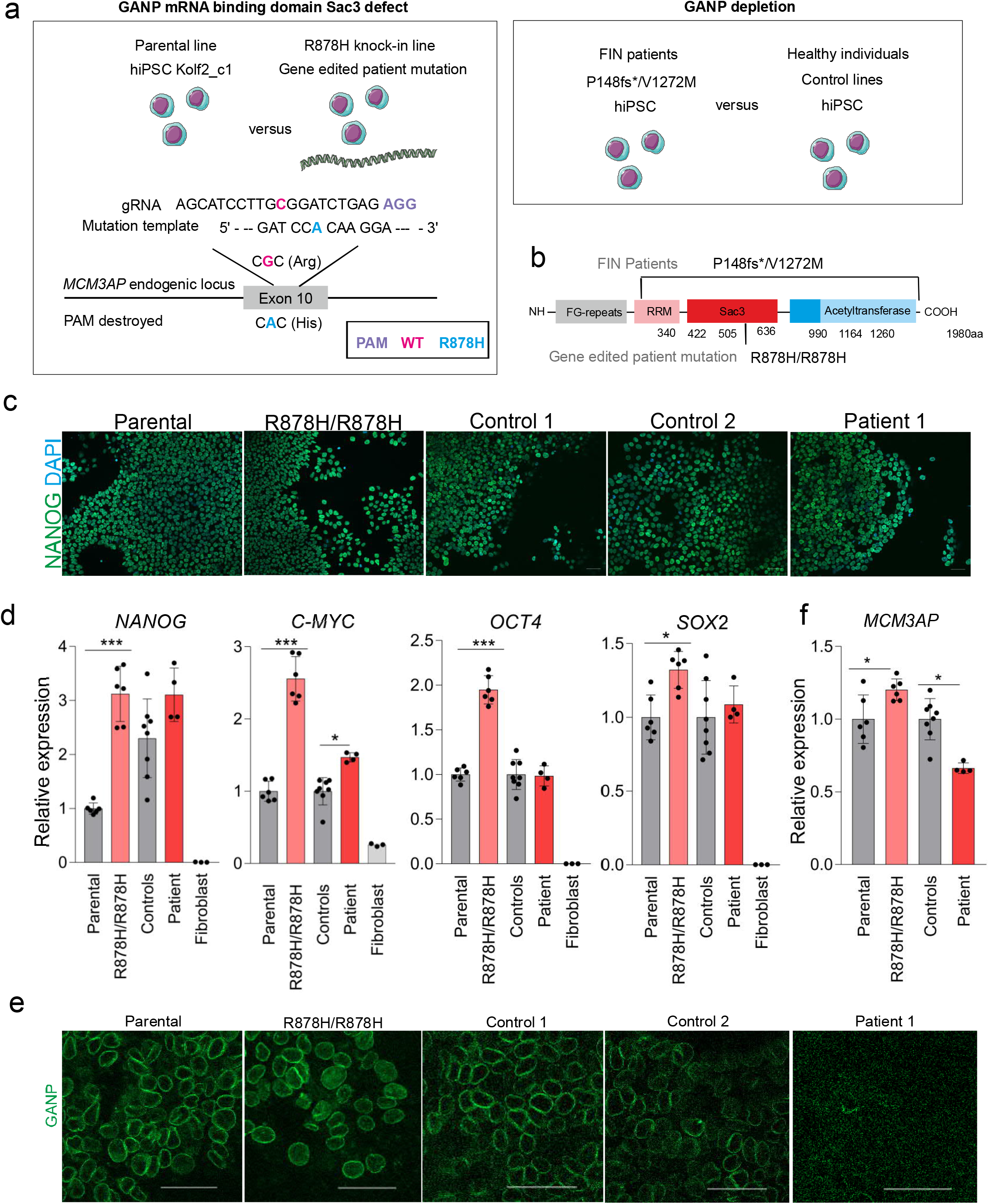
GANP-deficient hiPSC are pluripotent. a. Cell lines used in this study. Gene editing was used to generate the homozygous R878H Sac3 domain mutation in *MCM3AP* (GANP) on Kolf2_c1 parental hiPSC line, which was used as the isogenic control. In addition, fibroblasts from two FIN patients carrying P148fs*/V1272M were reprogrammed, and compared to healthy control hiPSC. b. Localization of the modelled disease variants in GANP domains. c. Immunocytochemistry to validate pluripotency with Nanog. DAPI indicates nuclei. Scale bar 50 µm. d. qRT-PCR of pluripotency markers *OCT4, NANOG, C-MYC* and *SOX2* relative to *GAPDH* expression (n=4-6). Fibroblasts are shown as a negative control. Patient values are relative to controls’ and R878H/R878H to parental average. e. Immunocytochemistry of GANP in hiPSC lines. Scale bar 50 µm. f. qRT-PCR of *MCM3AP* expression relative to *GAPDH* (n=3-6) in hiPSC. * p<0.05, *** p<0.0001.

We also reprogrammed skin fibroblasts of patients FIN1 and FIN2, who carry V1272M and P148*fs48 compound heterozygous variants, which result in depletion of GANP [61,63], and of two healthy unrelated controls (Figure 1A,B).

Pluripotency of all the above mentioned hiPSC was confirmed with Nanog immunocytochemistry (Figure 1C) and by qRT-PCR of pluripotency genes (Figure 1D). Tested pluripotency genes had higher expression in edited cells compared to parental hiPSC (*NANOG, OCT4*, and *C-MYC* p<0.0001; *SOX2* p<0.05). For pluripotency gene expression, we also investigated two additional edited hiPSC clones with heterozygous R878H mutation in one allele and an indel in the other allele (clones 138 and 245, Supplementary Figure 1), and found increased pluripotency gene expression also in those (*NANOG*, p<0.0001). In patient-derived hiPSC, pluripotency genes were similarly expressed as in control hiPSC, apart from *C-MYC*, which was higher in the patient-derived cells (p=0.0060) (Figure 1D).

### Mutation type affects GANP levels in hiPSC

We have previously shown in patient fibroblasts that homozygous missense variants affecting the Sac3 domain of GANP do not alter *MCM3AP* mRNA or GANP protein levels, whereas compound heterozygous variants outside the Sac3 domain typically lead to lowered *MCM3AP* mRNA level and depletion of GANP at the nuclear membrane [61]. Here, we used immunostaining to detect GANP in the hiPSC lines. We found that GANP localized to nuclear pores in the R878H/R878H knock-in, parental and healthy control hiPSC lines, but was reduced in hiPSC of patient FIN1 (Figure 1E), in line with our previous findings in fibroblasts. *MCM3AP* mRNA levels were reduced by approximately 30% in patient hiPSC compared to healthy controls, whereas we observed a 25% increase in the R878H/R878H knock-in line compared to the parental line (Figure 1F). These results are consistent with the two types of disease-causing *MCM3AP* variants having different effects on the level of GANP at the nuclear envelope.

### Patient-derived and knock-in hiPSC differentiate into motor neurons

To determine the effects of GANP variants in a disease-relevant cell type, we next differentiated hiPSC lines into motor neurons [15] (Figure 2A). Differentiation into a neuronal lineage was validated by qRT-PCR analysis of the cytoskeletal markers neurofilament heavy (*NEFH*) and βIII-Tubulin (*TUBB3*), and into a motor neuron lineage by analysis of choline acetyltransferase (*CHAT*) and ISL LIM homeobox 1 (*ISL1*) (Figure 2B). Immunofluorescence analysis showed that differentiated cells from all hiPSC lines were positive for neurofilament medium (NEFM), ISL1, and homeobox gene Hb9 (Figures 1C,D). ISL1 positivity was above 92% by the end of differentiation. In essence, we found that all hiPSCs differentiated similarly into motor neurons.

**Figure 2.**
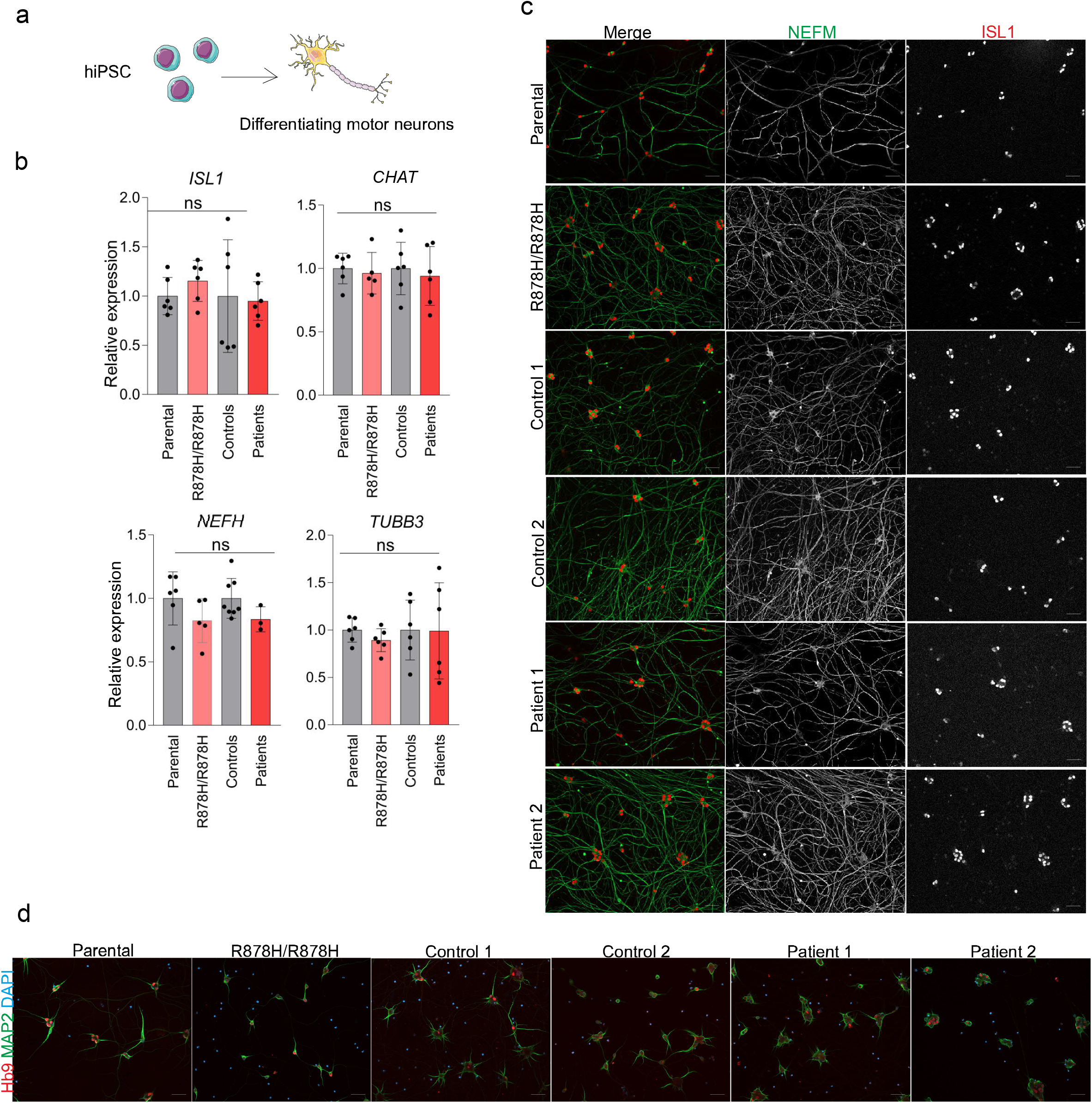
GANP-deficient hiPSC differentiate normally into motor neurons. a. hiPSC lines were differentiated into motor neurons. b. Gene expression validation of motor neuron transcripts *CHAT* and *ISL1* and neural transcript *NEFH* and *TUBB3* relative to *GAPDH* by qRT-PCR on day 23 of motor neuron differentiation (n=6 per clone). Patient lines compared to controls’ average and R878H/R878H to parental average. c. Immunocytochemistry of ISL1/2 (red) and NEFM (green) in neuronal cultures on day 23. d. Immunostaining of HB9 (red) and MAP2 (green). Scale bars 50 µm. ns = not significant.

### GANP is detected in patient-derived motor neurons

We immunostained the motor neurons and found that GANP localized to the nuclear pores in R878H/R878H knock-in, parental and healthy control motor neurons (Figure 3A). In contrast with our previous observations in FIN patients’ fibroblasts, in which GANP staining was depleted at the nuclear envelope, we detected GANP staining in patient-derived motor neurons. Nonetheless, also some abnormally staining nuclei were found particularly in FIN2 patient samples (Figure 3A). Patient-derived motor neurons had approximately 30% reduced *MCM3AP* mRNA levels in comparison to healthy controls (p=0.0007), whereas the levels were unchanged in R878H/R878H knock-in motor neurons (Figure 3B). Staining with NPC marker Mab414 appeared normal, suggesting that the nuclear envelope was intact in mutant and control motor neurons (Figure 3C).

**Figure 3.**
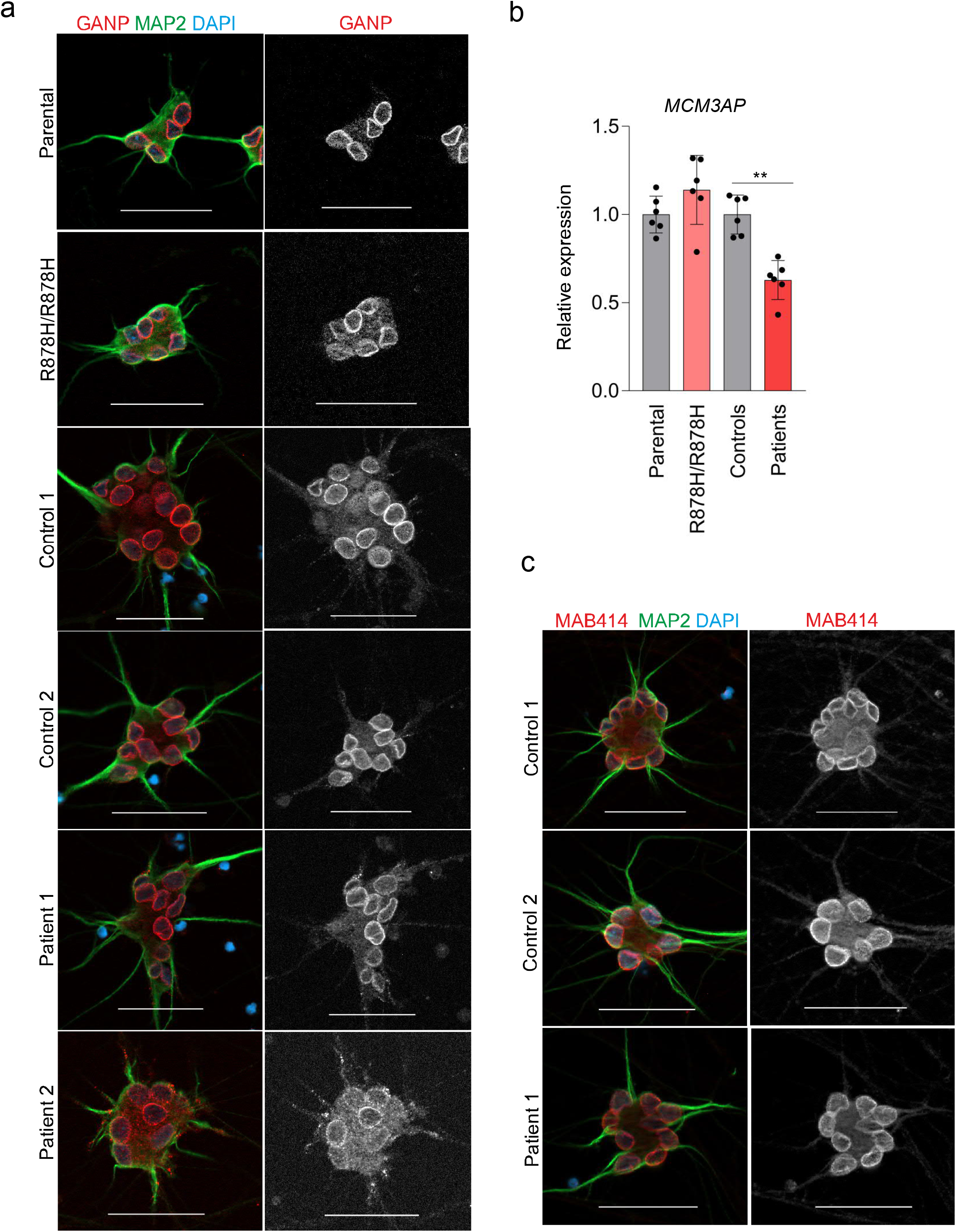
GANP localization and expression in human motor neurons. a. Immunocytochemistry of GANP (red) in MAP2 (green) positive neuronal cultures. DAPI stains for nuclei shown in blue. Scale bar 50 µm. b. Expression of *MCM3AP* in motor neurons analysed by qRT-PCR, relative to *GAPDH* (n=3-6 per clone). c. Immunocytochemistry of nuclear pore marker Mab414 (red) and neuronal marker MAP2 (green). DAPI is shown in blue. Scale bar 50 µm. ** p<0.001.

### GANP regulates gene expression in motor neurons

Given the roles of GANP in RNA processing, we next assessed whether mutant GANP affected gene expression in differentiated motor neurons by sequencing total RNA isolated from R878H/R878H knock-in neurons and isogenic control cells (Figure 4A). The motor neuron differentiation protocol we utilized yielded mostly homogeneous neuronal cultures, although as differentiation progressed, mitotic progenitors were also occasionally seen. For RNA sequencing we thus collected the neurons on day 22 of differentiation, and only from wells which did not have progenitor cells.

**Figure 4.**
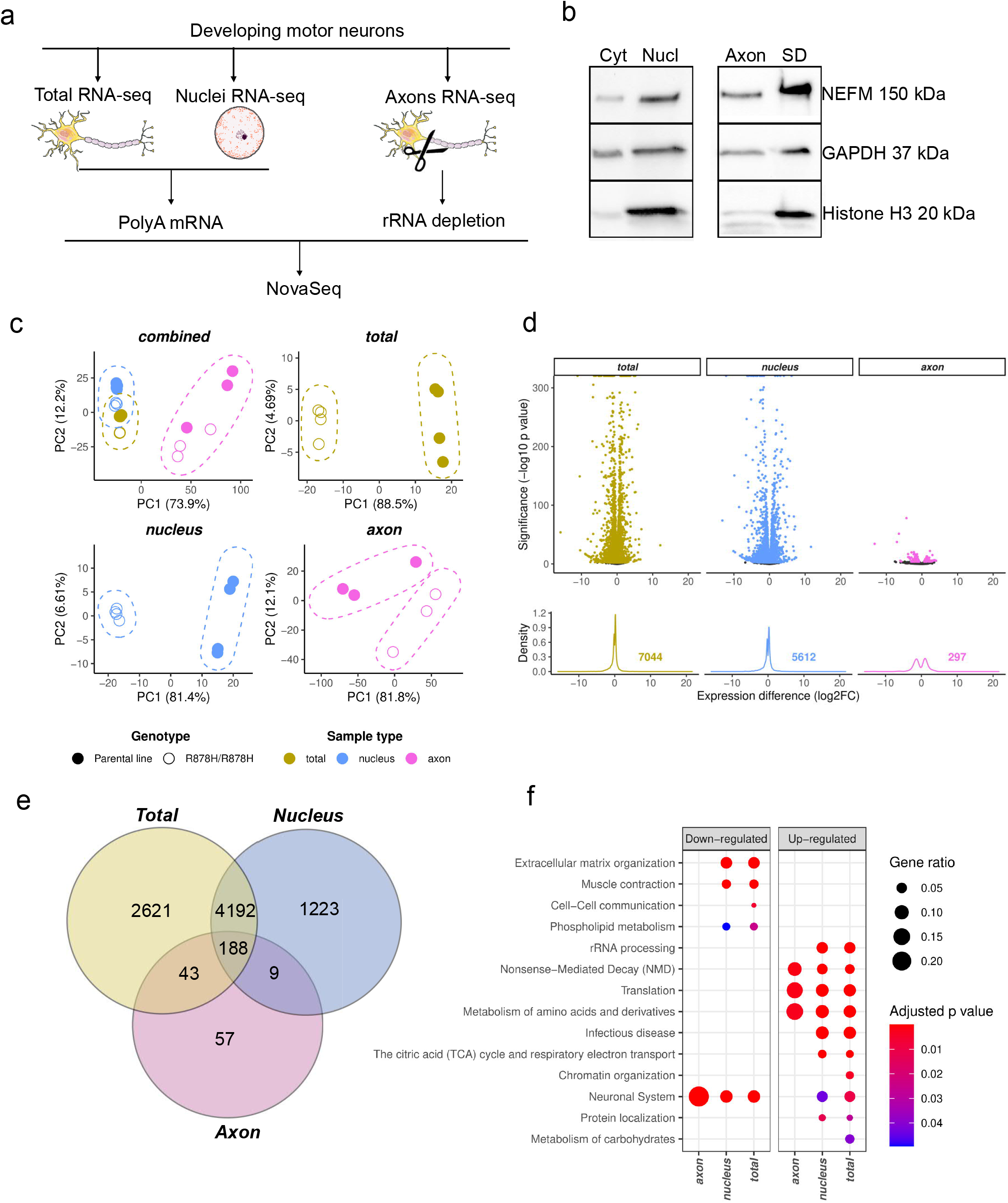
GANP is a major regulator of gene expression in human motor neurons. a. Schematic of workflow used for RNA sequencing experiments. b. Validation of nuclear and axonal isolations by western blotting of histone H3 (20 kDa), GAPDH (37 kDa), and NEFM (150 kDa). SD=somatodendritic fraction. c. Principal component analysis of RNA sequencing datasets. d. Density plot of differentially expressed genes in total neuron population and nuclear and axonal fractions. e. Venn diagram of the differentially expressed genes across the three different RNA sequencing datasets. f. Gene-set enrichment analysis of altered Reactome pathways.

Since GANP has roles in the nuclear export of RNA we also performed RNA sequencing on separate nuclear and axonal RNA fractions (Figure 4A). For the nuclear subcompartment, validation of the purity of the isolated nuclear fraction by western blotting showed highly enriched histone H3 protein (Figure 4B). To analyse transcripts specific to axons, we cultured motor neurons in Xona Chip devices consisting of microfluidic axon isolation technology. We then dissociated motor neurons from porous 1 µm filter inserts to obtain an isolated axon compartment [4]. The coating of the filter bottom with laminin encourages the attachment of axons, whilst the somatodendritic compartment grows on top of the filter. We validated the purity of our isolation of axons by western blotting, which showed low histone H3 levels (Figure 4B). To collect sufficient RNA for axon-Seq, samples from 6 separate wells for each replicate were combined, and in addition ribosomal RNA was depleted.

Principal component analysis of the RNA sequencing data showed that the sample types (total, nuclear, and axonal) clustered away from each other, indicating successful subcompartment isolation. Axon samples separated more from total and nuclear samples, demonstrating the uniqueness of the axonal transcriptome [38]. The overall motor neuron axon transcriptome was similar to previously reported datasets demonstrating enrichment in mtDNA- and nuclear-encoded mitochondrial respiratory chain genes, neurofilaments and expression of *TMSB10, YBX1* and *STMN2* [27,36,38]. Samples of the same genotype clustered together based on their gene expression, particularly for total and nuclear RNA, whereas axon samples showed more variation (Figure 4C).

Total RNA sequencing of R878H/R878H knock-in motor neurons showed that GANP has a major role in gene expression regulation in human motor neurons. Out of the 18,146 genes that we detected to be expressed in motor neurons, 7,044 (∼39%) were differentially expressed (DE) between R878H/R878H and parental motor neurons (adjusted p-value <0.01) (Figure 4D). The results for nuclear RNA were similar, showing 5,612 DE genes. More than 50% of the DE genes were the same in total and nuclear RNA (Figure 4E). Axon-Seq, by contrast, showed only 297 altered genes.

Overall pathway analysis of DE genes by Reactome enrichment indicated downregulation of genes related to neuronal system and extracellular matrix organization (Figure 4F). Several crucial axon guidance and maintenance genes were downregulated in GANP-deficient motor neurons including *SLITRK1, SLITRK3, DRAXIN, LSAMP* and axon repulsive genes involved in glutamatergic synapses (*NETO1, NETO2, SHISA9, GRIN1, NPTX1*) and neuronal system genes (*GABRG2, SYN2, SYT9)* (Figure 5).

**Figure 5.**
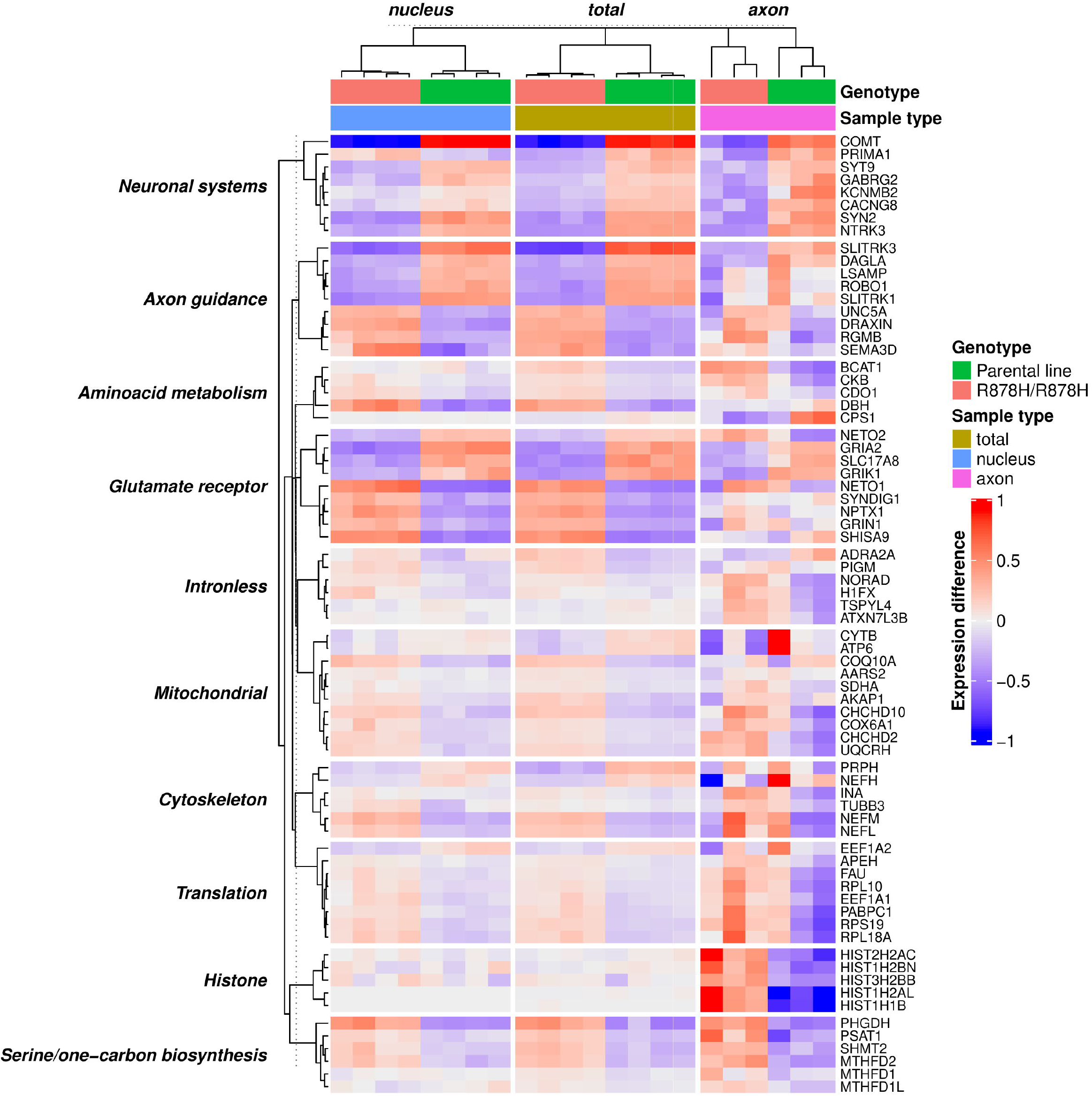
Heatmap of selected differentially expressed genes in GANP-deficient motor neurons. The heatmap shows representative genes from multiple enriched Reactome categories. To highlight differences among samples, the expression values (log1p FPKM) were scaled to gene averages separately for each sample type (total, nucleus and axon).

The overall upregulated genes in GANP-deficient motor neurons were associated with pathways involved in mitochondrial respiration, and in protein synthesis such as ribosomal subunits and translation elongation. This could indicate compensatory effects in response to altered mRNA export from the nucleus (Figure 4F, Figure 5). Interestingly, mtDNA-encoded transcripts were downregulated, whereas nuclear-encoded mitochondrial transcripts were upregulated, suggesting a nuclear response to mitochondrial dysfunction. Related to energy metabolic changes, CKB (brain-type creatine kinase) involved in compartmentalized ATP production and consumption was also upregulated [47]. Furthermore, upregulation of the serine and one-carbon biosynthesis pathway (*PHGDH, PSAT1, SHMT2, MTHFD2*) was detected, which is commonly observed in mitochondrial defects and as part of the integrated stress response [11,16].

The axonal transcriptome of GANP mutant motor neurons showed decreased levels of transcripts related to synaptic function such as glutamate ionotropic receptor subunits (Figure 5). Increased levels of a number of histone genes (for example *HIST2H2AC, HIST1H2BN, HIST1H1B)* were observed in mutant axons (Figure 5). As histone mRNAs are not expected to be translated in axons, this finding may have resulted from enrichment of histone mRNAs lacking polyA tails. However, we previously found in patient fibroblasts that GANP may specifically influence the export of intronless gene mRNAs [61], and it is possible that histone genes, which are intronless, escape the regulation by mutant GANP in neurons. Indeed, we identified a subset of intronless genes that were differentially expressed in motor neurons and in the nuclear subcompartment *(ADRA2A, PIGM, NORAD, ATXN7L3B)*.

Although genes related to neuronal functions were generally downregulated in knock-in motor neurons, neurofilament genes *NEFL* and *NEFM* were increased (p<0.0001) (Figure 5). Interestingly, by using qRT-PCR we found that *NEFM* expression was also increased in patient-derived motor neurons compared to controls (Figure 6A). The upregulation of neurofilament genes may be linked to our previous observation that *NEFL* expression is regulated by changes in protein synthesis [45]. Another possibility is that neurofilament gene expression was increased as a response to axonal injury. Indeed, neurofilaments are released by neurons in neurodegenerative diseases [25]. We therefore measured NEFL protein in the culture media but found that GANP knock-in or patient-derived motor neurons did not release more NEFL than control cells (Figure 6B). We also used microfluidic chips to test if the altered neurofilament gene expression affected the regeneration capacity following axotomy, but we did not detect defective axon regrowth in GANP knock-in motor neurons (Figure 6C).

**Figure 6.**
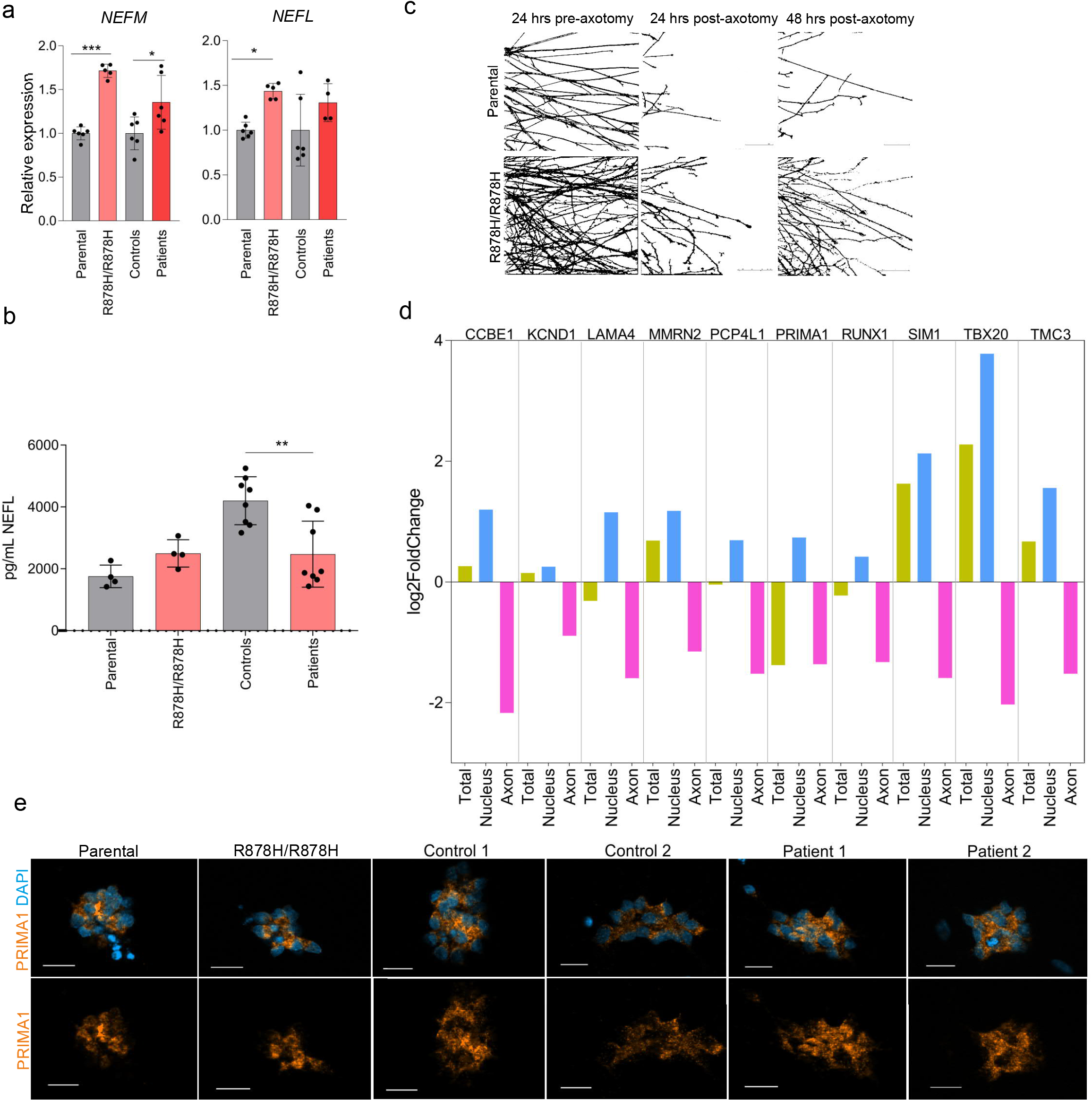
Nuclear mRNA retention in GANP-deficient motor neurons. a. mRNA expression of *NEFL* and *NEFM* in motor neurons by qRT-PCR, relative to *GAPDH*. b. NEFL measurement from neuronal culture medium. c. Axon regeneration assay. Images shown from 24 hrs before axotomy, immediately following axotomy and 24/48 hrs post-axotomy. One device analysed per line. Scale bar 100 µm. c. Quantification of axon thickness from light microscopy images 24 hrs before axotomy. d. Fold changes of genes that were upregulated in nuclear fraction and downregulated in axons, suggestive of nuclear mRNA retention e. RNA *in situ* hybridization of *PRIMA1* mRNA foci in motor neurons (orange). DAPI (blue) stains the nuclei. * p<0.05, ** p<0.001, ***p<0.0001.

We hypothesized that a defect in selective mRNA export should lead to nuclear mRNA retention and thus axonal depletion of a set of mRNAs. Thus, we analysed the RNASeq data for transcripts that were upregulated in GANP knock-in nuclei and downregulated in axons. However, only few such genes were detected (Figure 6D). One of those was *PRIMA1*, involved in organizing and anchoring acetylcholinesterase at neural cell membranes [41]. We studied the localization of *PRIMA1* mRNA by RNAScope but did not observe any obvious nuclear retention in the GANP mutant neurons (Figure 6E). Thus, we conclude that the R878H Sac3 mutation in GANP does not cause large-scale nuclear retention of mRNAs in human motor neurons but has a major effect on motor neuronal gene regulation.

### GANP regulates gene splicing in motor neurons

Finally, we analysed the effects of the R878H mutation on mRNA splicing in human motor neurons by KissSplice analysis. We observed a large number of significant alterations in splicing in motor neurons, in 421 genes, and fivefold more (in 2224 genes) in nuclear RNA sequencing (Figure 7A). The majority of these involved the usage of alternative splice sites, or were exon skipping or intron retention events. Splicing changes were observed both in genes that had an overall decreased mRNA expression, and in genes that were not differentially expressed (Figure 7B,C). As an example, a higher percentage of exon skipping was detected for *GABRG2*, inhibitory gamma-aminobutyric acid receptor subunit, which is alternatively regulated in neuronal development [56] (Figure 7D).

**Figure 7.**
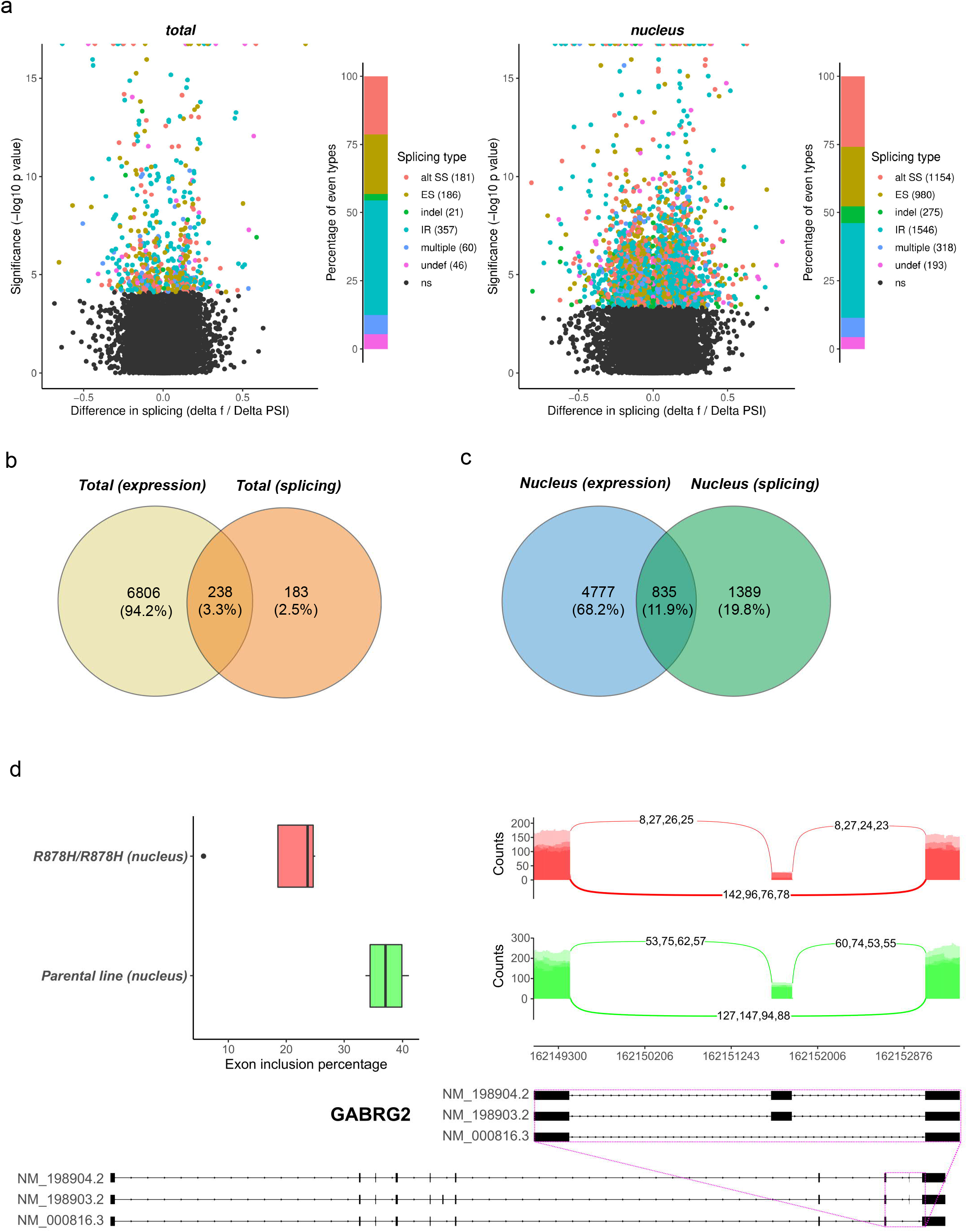
GANP regulates gene splicing in motor neurons. a. Volcano plot of splicing changes and types in total and nuclear RNA sequencing. b. Venn diagram indicating the numbers of differentially expressed and spliced genes in total RNA sequencing. c. Venn diagram indicating the numbers of differentially expressed and spliced genes in the nuclear compartment. d. Box plot indicating exon inclusion percentages in *GABRG2*. Sashimi plot around alternatively spliced exon. The values on the lines indicate the read counts for junction reads (reads that maps partially to more than one exon). The location of the splicing event is indicated with the refseq models.

## DISCUSSION

GANP was initially identified as an abundant protein in splenic germinal centers, which are responsible for development of B cells [30], and subsequently the role of GANP has been extensively characterized in the context of B cell affinity maturation [46]. The recognition of GANP as a mammalian mRNA export factor [59] was preceded by the identification of the yeast homologue of GANP, Sac3 [9,12,42], and the Drosophila homologue Xmas-2 [29]. Depletion of both Sac3 and Xmas-2 results in nuclear poly(A)+ RNA accumulation. Later studies showed that GANP stably associates with the nuclear pore basket [54], forming a scaffold for ENY2, PCID2 and centrin binding at TREX-2, thus firmly linking GANP to mRNA export [20]. Interestingly, GANP was then shown in HCT116 colon carcinoma cells to promote the export of highly expressed and short-lived transcripts that function particularly in RNA synthesis and processing, indicating that GANP can enable rapid changes in gene expression [58].

The identification of *MCM3AP* as a disease-linked gene causing a peripheral neuropathy syndrome prompted us to study GANP’s role in a disease relevant cell type. Although the phenotypes associated with GANP mutations are not restricted to peripheral nerves, motor neurons, which have a requirement for mRNAs to be transported extremely long distances into axon terminals, are particularly affected. Pathogenic variants in the mRNA export factor GLE1 also underlie a motor neuron disease, suggesting that motor neurons are vulnerable to inefficient mRNA export [10,21,39]. Reduced mRNA availability could have a major effect on local protein synthesis in axons, which is now recognized to be important for axon maintenance and synaptic function [34].

We aimed to identify human motor neuron specific genes that are regulated by GANP by utilizing a neuropathy-causing Sac3 domain mutant, which we generated by CRISPR/Cas9 gene editing in hiPSC. The homozygous missense variant R878H does not change the localization or the amount of GANP at the nuclear envelope but introduces local structural changes in the winged helix region of Sac3, thus impairing RNA binding [61]. We previously showed that GANP variants can be divided into two groups – compound heterozygous variants leading to GANP depletion, and homozygous Sac3 domain missense variants. Both cause loss-of-function, but notably the Sac3 variants associate with milder motor phenotypes than the GANP depletion variants [61-63]. This suggests that R878H and other Sac3 mutants cause partial impairment of GANP function. Unexpectedly, in the patient-derived motor neurons GANP was not strongly depleted, in contrast to what we observed in fibroblasts and iPSC from the same individuals [61]. The reason for this discrepancy is not known, but since the compound heterozygous mutations carried by these individuals include a frameshift variant in the first exon of *MCM3AP*, we speculate that the downregulation of nonsense-mediated decay response, which has been shown to be required for neuronal differentiation [19], may have resulted in higher amount of residual GANP.

Our results showed that motor neurons deficient for the RNA binding function of GANP differentiated normally but had major changes in gene regulation, including abnormal gene expression and splicing. Mutant GANP deregulated a large number of genes required for motor neuron functionality, whereas transcripts required for protein synthesis were upregulated, which we suggest is a compensatory response to abnormal mRNA availability. Expression of nuclear-encoded mitochondrial genes was also upregulated, which may compensate for the reduced availability of those transcripts in axonal protein synthesis that is needed to maintain functional mitochondria. Mitochondrial transcripts are among the most abundant mRNAs in axons [2,5,14,40,52,60], indicating their importance in local maintenance of axonal and synaptic mitochondria. We also observed a transcriptional upregulation of the serine and one carbon biosynthesis, an early-stage of integrated stress response, which may be caused by mitochondrial dysfunction [7,11,16]. These neuronal compensatory responses may be of relevance in patients who have the permanent impairment of GANP but have not been observed in studies of transient GANP depletion [1,58].

Previous studies of GANP depletion in diverse non-neuronal cell types have resulted in expression changes in genes that were largely distinct from those we detected here in GANP mutant motor neurons. However, one of the rare common findings in all studies is the downregulation of the intronless gene *ATXN7L3B*, encoding an adaptor protein in the TREX-2 complex. It was downregulated by GANP siRNA knockdown in human colon carcinoma cells [58], in our study of patient fibroblast gene expression [61], in a recent study of a degron system induced GANP knockdown in a colorectal cancer line [1], and in the GANP mutant motor neurons. Together, these findings suggest that *ATXN7L3B* is a specific target of GANP regulation regardless of cell type. Interestingly, *ATXN7L3B* is also a subunit of the Spt-Ada-Gcn5 acetyltransferase (SAGA) complex, involved in histone modifications and thus in gene expression regulation [28]. The multiple effects that GANP has on gene regulation may thus be mediated by the SAGA complex, or through the interactions of GANP with the other large complexes that are interconnected from transcription to splicing and mRNA export.

In conclusion, we have shown that GANP is a major gene regulator in human motor neurons, with effects beyond its role in mRNA export. Large-scale nuclear mRNA retention was not observed in our neuronal model with the GANP Sac3 missense mutation, which suggests that the disease-associated impairment is moderate enough to enable the export of most mRNAs, but nevertheless limits mRNA availability, resulting in compensatory responses in protein synthesis and metabolism. Our findings are consistent with GANP-associated disease effects on projection neurons with long axons in human patients.

## MATERIALS AND METHODS

### hiPSC culture and maintenance

hiPSC were expanded and grown for experiments in E8/E8 Flex basal media with 50X supplement (Thermo) and Primocin (1:500). Cells were routinely passaged with Versene (Thermo) or EDTA and grown on Vitronectin (Thermo). When required, Revitacell/Y-27632 was added (1:1000). All cells used in this study were culture at 37°C, in a humidified atmosphere, normoxia and 5% CO2.

### Reprogramming

Human skin fibroblasts from two healthy controls and from two patients (P148fs*/V1272M) [63] were reprogrammed into pluripotent stem cells at Biomedicum Stem Cell Center (University of Helsinki, Finland). Patient cells were reprogrammed in oriP/EBNA1 backbone and expression of OCT3/4, SOX2, KLF4, MYC, LIN28 and shRNA against p53. All cell lines were validated for cell growth, hiPSC morphology and pluripotency gene expression by immunocytochemistry, RT-PCR and qPCR. hiPSC were cultured in Matrigel-coated (Corning) plates with E8-medium (Gibco) supplemented with E8-supplement (Gibco). Cells were passaged when confluent with 0.5 mM EDTA (Invitrogen) in phosphate-buffer saline (PBS).

### Gene editing

Gene editing was done following a previously published protocol [3]. Kolf2_C1 p.13 human induced pluripotent stem cells (hiPSCs) characterised by the HipSci consortium [32] were cultured in a humidified incubator at 37°C and 5% CO_2_. The cells were cultured in feeder-free conditions in 10 cm culture dishes in TeSR-E8 (Stem Cell Tech) medium on SyntheMAX (2µg/cm^2^) (Corning #3535). For passaging cells were washed with PBS (no Mg/Ca^2^), incubated with Gentle Cell Dissociation Reagent (Stem Cell Tech) at 37°C for 3 min and added TesR-E8 media.

The synthetic guide RNA (sgRNA) harbouring the R878H mutation (clones R878H/R878H and R878H/indel-138), AGCATCCTTGCGGATCTGAG, was ordered from Synthego following design of effective guides (https://wge.stemcell.sanger.ac.uk). A single stranded oligonucleotide(ssODN), sequence: 5-GCTGTGTGCTCACCGTGTACGCAAAGTTGAGCGCCCGGAGAGCATCCTTGTGGATCTGAGAGGAGGAGCGAAATCACTGCAGTCTCAGACGAAGGCCCAGT-3, was ordered from IDT. sgRNA GATCCGCAAGGATGCTCTCC GGG (reversed) and ssODN 5-ACTGGGCCTTCGTCTGAGACTGCAGTGATTTCGCTCCTCCTCTCAGATCCACAAGGATGCTCTCCGGGCGCTCAACTTTGCGTACACGGTGAGCACACAGC-3 was used to produce clone: R878H/indel-245. Desalted ssODN (IDT, Ultramer DNA oligo) had equal homology arms of 50 bases around the mutation.

Confluent 10 cm petri dish was dissociated with Accutase into single cells. 1 million cells were used for electroporation. *Sp*Cas9 ribonucleoprotein complexes (RNP) were formed with sgRNA and ssODN. sgRNA oligos were suspended in IDT duplex buffer solution at concentration 200µm and 500 pmol ssODN templates at final concentration of 100µm. Cas9 (purified from *E*.*coli*) suspended in Cas9 storage buffer and RNP was reconstituted to a final concentration of 4µg/µl. Cells were electroporated with Amaxa 4D using (Lonza #AAF-1002B, #AAF-1002x) suspended in P3 Primary cell 4D-Nucleofector X Kit L (Lonza, V4XP-3012) and CA137 program. Final concentration of Cas9 per 1 M cells was 20 µg and 20 µg sgRNA. Cells were processed and plated in TeSR-E8 media with Rock inhibitor (Y-27632) on SyntheMAX coated (5µg/cm^2^) 6-well plates. Cells were fed daily until 70% confluent. Recovered cells were dissociated three days later from semi-confluent wells with Gentle Cell Dissociation Reagent and frozen in 90% knock-out serum replacement with 10% DMSO.

### Karyotyping

To process cells for karyotyping, hiPSC were grown on 6 cm dishes until semi-confluent. Colecimid 10 µg/mL was added to the cells and placed in the incubator for 4 hrs. After two washes with PBS, the cells were dissociated with Tryple Select (Gibco). DMEM was added and cells were centrifuged 200 g for 5 min. The cells were incubated in KCl at +37°C. Cells were fixed with 25% Acetic acid in methanol and centrifuged in 200 g 5 min, and this was repeated in total three times. Karyotype analyses based on chromosomal G-banding were performed at Analisis Mediques Barcelona.

### Motor neuron differentiation

To differentiate hiPSC into motor neurons, we used a previous described protocol [55] with small modifications. In summary, confluent hiPSC were dissociated into low-attachment flasks with 0.5 mM EDTA and cultured in Neuronal basal medium (DMEM/F12/Glutamax, Neurobasal vol:vol, with N2 (Life Technologies), B27 (Life technologies), L-ascorbic acid 0.1 mM (Santa Cruz) and Primocin supplemented with 5 µM Y-27632 (Selleckchem)/40 μM SB43154 (Merck)/ 0.2 μM LDN-193189 (Merck/Sigma)/3 µM CHIR99021 (Selleckchem) for 2 days. Following five days the cells were cultured in basal media and 0.1 µM retinoic acid (Fisher)/0.5 µM SAG (Calbiochem) and for the next 7 days, BDNF and GDNF (10 ng/ml, Peprotech) were added. 20 µM DAPT (Calbiochem) was added for days 9-17. Day 10 neurons grown in spheroids were dissociated with Accumax (Invitrogen) on poly-D-lysine 50 µg/ml (Merck Millipore) and laminin 10 µg/ml (SigmaAldrich). Day 17 onwards until analysis, neurons were kept in BDNF, GDNF and CNTF. Media were changed every 2-3 days by replacing half of the medium. Developing motor neurons were analysed for experiments from day 22 onwards.

### cDNA synthesis and RT-PCR

iPSC and neuron total RNA was isolated with Nucleospin RNA kit (#740955.50, Macherey-Nagel). cDNA was synthesized according to manufacturer’s instructions with Maxima First Strand cDNA Synthesis Kit (#K1671, Thermo Fisher Scientific).

### qRT-PCR

qPCR was run on a CFX96 Touch Real-Time PCR detection system (Bio-Rad) with SYBR Green qPCR Kit (#F415S, Thermo Fisher Scientific) on a 96-well plate with gene-specific primers 10 µM and H_2_0. The cycling protocol was 95°C for 7 min, then 40 cycles of 95°C for 10 s and 60°C for 30 s. Normalization was done with *GAPDH* as a housekeeping gene using ΔΔcT method. Minimum of 3-4 technical samples were used for all studies. 12.5 ng of cDNA was used for hiPSC and 6.25 ng for neurons.

### Immunocytochemistry

Cells were washed three times with PBS, fixed with 4% paraformaldehyde for 15 min followed by three PBS washes. Cells were then permeabilized in 0.2% Triton X-100/PBS for 15 min at RT. Coverslips were washed three times in PBST (0.1% Tween-20) followed by block at RT with 5% BSA/PBST (#001-000-162, Jackson Immuno Research) for two hours. Primary antibody in 5% BSA/PBST was incubated overnight at 4°C. Following overnight incubation, coverslips were washed with PBST for 15 min three times, followed by incubation in secondary antibody in 5% BSA/PBST for one hour. The cells were finally washed three times for 10 min and coverslips plated on glass with Vectashield containing DAPI (#H-1200, Vector Laboratories).

### Antibodies

Antibodies were used at the following dilutions: 1:200 α-rabbit N-terminal GANP antibody for immunocytochemistry was (#198173, Abcam) with secondary antibody 1:1000 (488 goat-anti-rabbit #A11008, Alexa or 594 goat-anti-rabbit #110012, Alexa). To detect α-rabbit Nanog 1:200 (#4903, Cell Signalling), secondary 488 goat-anti-rabbit 1:1000 (A11008, Alexa) was used. Neuronal antibodies used for immunocytochemistry used were: 1:200 mouse α-Hb9 (#81.5C10-C, DSHB) with secondary antibody 1:1000: (#A11007, Alexa), 1:200 chicken α-MAP2 antibody (#AB5543, Millipore) with secondary antibody 1:1000 (#SA5-10070, Thermo). 1:1000 rabbit α-histone H3 (#4499, Cell Signalling) with secondary antibody 1:1000: 1:700 rabbit α-NEFM antibody for immunocytochemistry and 1:1000 for Western blotting (#20664-1-AP, Proteintech), with secondary antibodies 1:1000 (A11008, Alexa) and (#111-035-144, Jackson), respectively. To visualize mouse α-Isl1 1:200 (#39-4D5, DSHB), secondary 594 goat-anti-mouse was used 1:1000 (#A11007, Alexa). To visualize nuclear pores, mouse α-Mab414 1:200 (#ab24609, Abcam) was used with secondary antibody 594 goat-anti-mouse 1:1000 (#A11007, Alexa). For Western blotting, rabbit α-GAPDH 1:1000 (#14C10, Cell Signalling) with secondary 1:5000 (#111-035-144, Jackson) and mouse α-NEFL 1:1000 (#sc39073, Santa Cruz) with secondary 1:5000 (#115-035-146, Jackson).

### Axon isolation

Dissociated Accutase passaged single neurons d11 (400K) were plated in motor neuron differentiation media in 6 well hanging inserts with Boyden chamber device (1 μm pore, Falcon Cell Culture Inserts, #10289270, Thermo) for axon isolation (d22) similar to recent protocols [4,18]. Transmembrane filters were coated on both sides with poly-d-lysine and only on bottom side with laminin. Somatodendritic compartment was removed by scraping with PBS. The filter top was then washed 10 times with PBS RT strongly pipetting and then filter containing axons on bottom side was cut from the insert and washed 1 time with cold PBS. For RNA, PBS was removed by aspiration after spinning down and continued to RNA extraction with Nucleospin RNA kit (#740955.50, Macherey-Nagel) (incubated in the first buffer for 15 min after vortexing). Six filters combined into same conical tube for one technical replicate for RNA sequencing.

For isolating protein for validation by immunoblotting, the filter was then placed on ice following the washing steps and lysed in 30 uL RIPA containing Halt protease inhibitor (#1861284, Thermo) was added. All samples were vortexed and incubated on ice for 15 min. The samples were then centrifuged for 10 min at 14 000 x g in a cold centrifuge and supernatant was removed for protein. Three wells combined into same conical tube. Protein concentration was measured with BCA (#23227, Thermo).

### Nuclei isolation for RNA and protein

Neurons d22 (500K) were washed once with PBS RT, lysed in homogenizing buffer (0.3 M sucrose, 1 mM EGTA, 5 mM MOPS, 5 mM KH2PO4, pH 7.4). Cells were scraped and an aliquot was removed for total cell extract for protein samples. Cells were run through 22 G syringe and needle 8 times, an aliquot was taken for trypan blue staining and cells were observed under microscope for lysis. Cells were centrifuged 15 minutes at 6500 rpm at 4°C. Supernatant formed the cytoplasmic extract and pellet the nuclear extract. RNA was isolated with Nucleospin RNA kit (#740955.50, Macherey-Nagel) and RNA concentrations were measured with Denovix and TapeStation. For protein, nuclear and total cell extracts were lysed in 1 x RIPA with protease inhibitor, incubated on ice 5 min, centrifuged 14 000 x g for 10 min, removed supernatant for protein.

### Protein extraction and Western blotting

The cells were isolated with 1X RIPA (#9806, Cell Signalling) for 5 min on ice, scraped and centrifuged at 14 000 x g for 10 min at 4°C. Protein concentrations were measured with BCA (#23227, Thermo). For SDS-PAGE analysis, 5 µg of protein was loaded per well on a 10% TGX 10-well 30µl/well gel (456-1033, Bio-Rad) for 1 hr. at 150V. Molecular size of bands was detected with Protein Standard (#161-0374, Bio-Rad) and proteins loaded together with 4x Laemmli loading buffer (#161-0747, Bio-Rad) in 2-mercaptoethanol (#1610710, Bio-Rad). Gel was transferred into a nitrocellulose membrane (#1704158, Bio-Rad) and transferred using Mixed Molecular Weight protocol in Trans-Blot Turbo (Bio-Rad). The blot was washed blocked in 5% milk/TBST for 1 hr in RT. For immunoblotting analysis, antibodies were incubated overnight in 4°C in 2,5% milk/BSA in TBST. The blot was washed three times for five min in TBST. Secondary HRP antibodies (1:5000) in 1% BSA/TBST was incubated for 1 hr at RT on a shaker and washed three times 10 min in TBST. The blot was visualized with ECL reaction (#K-12049-D50, Advansta). The images were obtained with a Chemidoc imager (Bio Rad) and quantified on Image Lab software (Bio-Rad). Results were visualized on Graph Pad Prism Software.

### RNA sequencing and bioinformatics

For total neuron population, RNA was isolated from 6-well plates of Accumax dissociated and cultured single neurons (500K) four independent wells grown in poly d lysine and laminin grown in culture for 22 days. Samples were sequenced and analysed for Kolf2_c1 parental wild type and R878H/R878H knock-in line. Cells were washed with PBS once and RNA isolated from lysed cells Nucleospin RNA kit (#740955.50, Macherey-Nagel) according to manufacturer’s instructions including rDNAse treatment. RNA concentrations were measured with Agilent TapeStation and Nanodrop, RNA integrity was checked on a gel with Agilent TapeStation.

The RNA-seq libraries were made with NEBNext Ultra II Directional (PolyA) kit and sequenced with NovaSeq SP 2×150bp (Illumina) at Biomedicum Functional Genomics Unit (FuGU). The raw data was filtered with cutadapt [37] to remove adapter sequences, ambiquous (N) and low quality bases (Phred score < 25). We also excluded read pairs that were too short (<25bp) after trimming. The filtered read pair were mapped to the human reference genome (GRCh38) with STAR aligner [6]. Gene expression was counted from read pairs mapping to exons using featureCount in Rsubreads [33]. Duplicates, chimeric and multimapping reads were excluded, as well as reads with low mapping score (MAPQ < 10). The read count data was analyzed with DESeq2 [35]. We analyzed the effect of R878H/R878H mutation in *MCM3AP* by comparing the mutant samples to the parental cell line, separately for each sample type (total RNA, nuclear RNA and axonal RNA). We removed genes with low expression levels from the analysis (<50 reads across all samples). PCA was calculated with prcomp using variance stabilizing transformation (vst) on the read counts. The differentially expressed genes (FDR<0.01) were analyzed for enrichment separately for the up- and down-regulated genes using clusterProfiler [64] against the Reactome pathways [8], KEGG [22], Gene ontology [53] and Disease ontology [48]. For background genes we used all the genes that were detected as expressed in the samples (above the low coverage cutoff, >50 reads). We analyzed alternative splicing dynamics with KisSplice [44]. The results from the gene expression analysis together with the raw sequences were deposited to GEO, under accession GSE162781.

### RNAScope

RNA *in situ* hybridization was performed on fixed adherent hiPSCs grown on coverslips using RNAscope Multiplex Fluorescent Reagent Kit V2 (#323100, Advanced Cell Diagnostics) for target detection according to the manual. Briefly, 4% paraformaldehyde fixed cells were dehydrated in increasing ethanol concentrations (50%, 70%), and stored in 100% EtOH at - 20°C. The day of RNA *in situ* hybridization, were cells rehydrated in decreasing concentrations of ethanol (70%, 50%) and in PBS. Then treated with hydrogen peroxide for 10 min at RT, washed with distilled water, and treated with 1:30 diluted protease III for 10 min at RT, and washed with PBS. Probes Hs PRIMA1 (ACD, #900601) were hybridized for 2 h at 40°C followed by signal amplification and developing the HRP channel with TSA Plus fluorophore Cyanine 3 (1:1500) (NEL744001KT, Perkin Elmer) according to the manual. Cells were counterstained with DAPI and mounted with ProLong Gold Antifade Mountant (#P36930, Invitrogen). Cells were imaged with Zeiss Axio Observer microscope (Zeiss, Germany) with a 20X air objective and Apotome.2.

### Axon regeneration assay

Neurons were plated and axons grown on microfluidic chips (#XC450, Xona microfluidics). To dissociate, axons were washed with media until they detached. Axon growth was visualized by light microscopy on days 21-23 of differentiation, before and after axotomy. Axon masks from images were generated with Fiji.

### Measurement of NEFL from culture medium

For measurement of NEFL, 1 mL of culture medium was collected from single 6-well neuronal cultures (4 technical replicates) at day 23 of neuronal differentiation. Samples were centrifuged at 13000 x g for 10 min in a cold centrifuge and supernatant frozen in −20°C. NEFL levels were quantified using the Quanterix single molecule array [43] (Simoa, Billerica, MA, USA) HD-1 analyzer and Quanterix Simoa NF-Light Advantage Kit (ref. 103186) according to manufacturer’s instructions. Briefly, frozen medium samples were slowly thawn on ice, mixed, and centrifuged at 10000g for 5 min at room temperature and diluted 1:50 in sample diluent prior to loading to a 96-well plate. Samples were measured in duplicate. All measured concentrations were within the calibration range and the mean coefficient of variation (CV) of sample replicates was 4.5%. A CV of <15% was considered acceptable between the replicates. The data are shown as mean of the replicate samples ± SD (pg/mL).

### Statistics

One-way ANOVA was used for comparing means between different groups and unpaired two-tailed t-test for comparisons of two groups. Significance was measured by p <0.05 in qRT-PCR and axon microfluidics experiments. Statistical tests were performed in Graph Pad Prism 7.04/8.03. All measured data points were included in the analysis.

## ACKNOWLEDGEMENTS

Riitta Lehtinen, Jana Pennonen and Anni Laitinen are thanked for technical assistance. We thank Tarja Kokkola, PhD, at UEF Biomarker Laboratory, Kuopio, Finland, for her kind help and advice on the Simoa-based medium NEFL measurements. We acknowledge Biomedicum Functional Genomics Unit of the Institute of Biotechnology, University of Helsinki, and Biocenter Finland Stem Cell Center. Funding: This work was supported by Doctoral Programme Brain & Mind (RW), UEF Doctoral Program in Molecular Medicine (NH), University of Helsinki (EY, HT), Academy of Finland (EY, HT, AH), Instrumentarium Science Foundation (RW), HUS Helsinki University Hospital (EY), Emil Aaltonen Foundation (EY) and Sigrid Juselius Foundation (HT).

## AUTHOR CONTRIBUTIONS

RW, EY, JS and HT conceived the ideas and planned the experiments. RW, MS, MTS, SH developed and modified the methodologies used. JK did the bioinformatics analysis and implemented the codes. RW, MS and NH completed and validated the experiments and did the formal analyses. NH, S-K.H and AH produced the NEFL data. SHä conducted RNAScope experiment which OC supervised. RT-M participated in AxonSeq experiment. RW gene edited hiPSC, which JS, MW and AB supervised. RW, MS, JK prepared the figures and visualization of the data. RW wrote the initial manuscript and HT and JS reviewed and edited the draft. All authors reviewed the final manuscript.

## SUPPLEMENTAL INFORMATION

**Supplemental table 1**.

Differential expression analysis results.

**Supplemental table 2**.

Gene-set enrichment analysis results.

**Supplemental table 3**.

Splicing analysis results.

**Supplemental table 4**.

Gene-set enrichment analysis results for splicing changes.

---

**Supplementary Figure 1.**
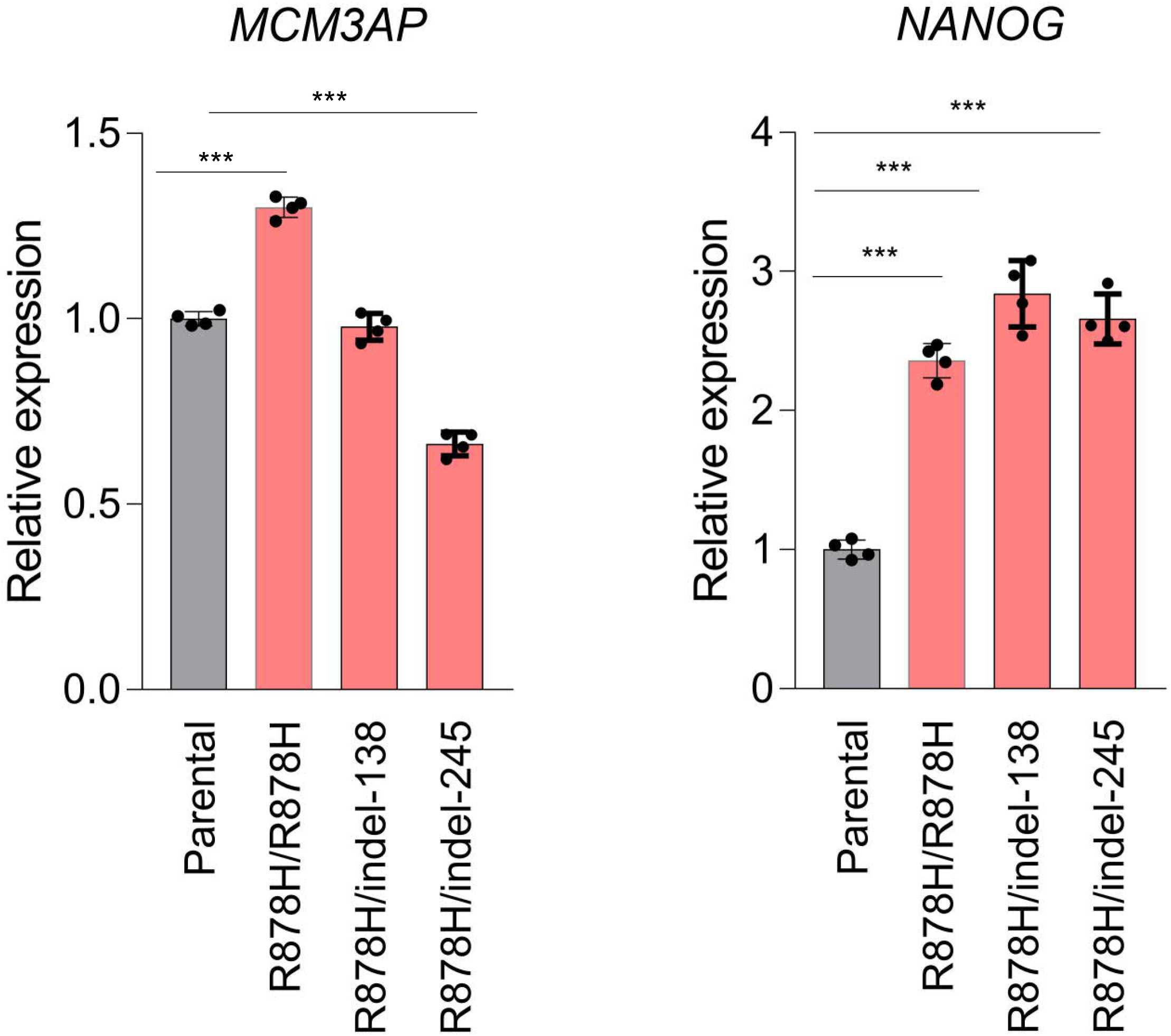
Increased expression of pluripotency marker *NANOG* in iPSC. qRT-PCR of *MCM3AP* and pluripotency marker *NANOG* in different gene-edited iPSC lines. * p<0.05, *** p<0.0001.

